# Preclinical evaluation of the efficacy and safety of AAV1-hOTOF in mice and non-human primates

**DOI:** 10.1101/2023.08.22.554252

**Authors:** Longlong Zhang, Hui Wang, Mengzhao Xun, Honghai Tang, Jinghan Wang, Jun Lv, Biyun Zhu, Yuxin Chen, Daqi Wang, Shaowei Hu, Ziwen Gao, Jianping Liu, Zheng-Yi Chen, Bing Chen, Huawei Li, Yilai Shu

**Affiliations:** ENT institute and Otorhinolaryngology Department of Eye & ENT Hospital, State Key Laboratory of Medical Neurobiology and MOE Frontiers Center for Brain Science, Fudan University, Shanghai, 200031, China; Institutes of Biomedical Science, Fudan University, Shanghai, 200032, China; NHC Key Laboratory of Hearing Medicine, Fudan University, Shanghai, 200031, China; Department of Otolaryngology-Head and Neck Surgery, Graduate Program in Speech and Hearing Bioscience and Technology and Program in Neuroscience, Harvard Medical School, Boston, MA 02115, USA; Eaton-Peabody Laboratory, Massachusetts Eye and Ear, 243 Charles St., Boston, MA 02114, USA

**Keywords:** Hereditary hearing loss, Gene therapy, Dual AAV delivery, DFNB9, AAV1-hOTOF, Efficacy and safety

## Abstract

Pathogenic mutations in the *OTOF* gene cause autosomal recessive hearing loss 9 (DFNB9), one of the most common forms of auditory neuropathy. There is no biological treatment for DFNB9. Here, we designed an *OTOF* gene therapy agent by dual AAV1 carrying human *OTOF* coding sequences with the expression driven by the hair cell-specific promoter *Myo15*, AAV1-hOTOF. To develop a clinical application of AAV1-hOTOF gene therapy, we evaluated its efficacy and safety in animal models by pharmacodynamics, behavior, and histopathology. AAV1-hOTOF inner ear delivery significantly improved hearing in *Otof* ^−/−^ mice without affecting normal hearing in wild-type mice. AAV1 was predominately distributed to the cochlea although it was detected in other organs such as the central nervous system and the liver, and no obvious toxic effects of AAV1-hOTOF were observed in mice. To further evaluate the safety of *Myo15* promoter-driven AAV1-transgene, AAV1-GFP was delivered into the inner ear of *Macaca fascicularis* via the round window membrane. AAV1-GFP transduced 60-94% of the inner hair cells along the cochlear turns. AAV1-GFP was detected in isolated organs and no significant adverse effects were detected. These results suggest that AAV1-hOTOF is well tolerated and effective in animals, providing critical support for its clinical translation.

## Introduction

Disabling hearing loss is a common sensory deficit, with about 466 million people affected worldwide, ^1^ and genetic defects account for 60% of congenital hearing loss.^2,3^ To date, clinical therapeutic strategies mainly rely on hearing aids or cochlear implantation to alleviate genetic hearing loss.^4–6^ However, these approaches are limited by their sensitivity and their ability to perceive music rhythm and natural sounds especially in noisy environments.^7–13^ Thus, there is an urgent need to explore novel treatment strategies for hereditary hearing loss, and with technical developments in the delivery and genome editing tools, etiology-based gene therapy has emerged as a promising strategy for hereditary hearing loss.^14–17^

DFNB9 is a form of congenital profound deafness due to mutations in the *OTOF* gene^15, 18, 19^ and accounts for 2-8% of hereditary deafness.^18–21^ Over 200 pathogenic mutations have been identified in the *OTOF* gene.^22, 23^ Pathogenic mutations in *OTOF* gene disrupt otoferlin function in the inner hair cells (IHCs),^24^ thereby leading to almost completely abolished exocytosis function of IHCs resulting in sensorineural hearing loss.^25, 26^

Due to the large size of *OTOF* cDNA (∼6 kb), gene replacement strategies based on the nucleic acid or protein recombination of dual adeno-associated virus (AAV) rescued hearing loss in the *Otof* ^−/−^ mouse models.^15–17^ However, these strategies still lack the safety evaluations required to advance to the clinical stage. In previous studies, the delivery tools (AAVs) used ubiquitous promoters such as CMV and CAG lacked target specificity for HCs in the cochlea,^15–17^ which may result in *OTOF* expression in non-hair cells, which may lead to adverse effects.^27, 28^ As a result, it is unclear whether the previous *OTOF* treatments pose potential safety risks. To minimize the potential risk, a cell-type specific promoter is required to restrict the *OTOF* expression to hair cells that are physiologically relevant.

We designed a treatment system – AAV1-hOTOF – that consists of the HC-specific promoter *Myo15* and the human *OTOF* coding sequences (CDS) mediated by AAV1, which has been approved to treat multiple human diseases.^29–33^ We report the assessment of the efficacy and safety of the AAV1-transgene driven by the *Myo15* promoter in mice and non-human primates through pharmacodynamics measurements, behavioral tests, and histopathology analysis. This study provides valuable efficacy and safety data for conducting clinical trials of AAV1-hOTOF.

## Results

### Hearing in *Otof* ^−/−^ mice was rescued after AAV1-hOTOF injection

To promote otoferlin gene therapy for clinical application, we first selected a relative safe AAV1 as delivery tool, and confirmed that AAV1 transfected IHCs effectively (**Figure S1A**-**B**), consistent with previous reports.^34–36^ Next, we screened out an ideal recombination strategy for *OTOF* gene therapy in nucleic acid level by western blot and auditory tests (**Figure S2A**-**C**), indicating that AK strategy outperformed other two strategies (TS and AP).^37^ For this, we used a dual-AAV1-Hyb(AK) approach in which the human *OTOF* CDS was driven by the HC-specific promoter *Myo15*, named AAV1-hOTOF.

To further systematically evaluate the efficacy of AAV1-hOTOF in treating *Otof* ^−/−^ mice, we first injected newborn mice via the round window membrane (RWM) at postnatal day 0-2 (P0-P2). As shown in **Figure 1A**, the untreated *Otof* ^−/−^ mice were profoundly deaf, with no identifiable auditory brainstem response (ABR) waves elicited by the maximum 90 dB sound pressure level (SPL), while the AAV1-hOTOF injected ear of *Otof* ^−/−^ mice displayed distinct ABR waves. In order to further explore the relationship between dose and efficacy, four doses were tested, including the high-dose group (6×10^10^ viral genomes (vg)/cochlea), the intermediate-high-dose group (3×10^10^ vg/cochlea), the intermediate-dose group (1.5×10^10^ vg/cochlea), and the low-dose group (7.5×10^9^ vg/cochlea). ABRs in response to click and tone-burst stimuli were recorded 1 month after injection. In the injected ears, the click ABR thresholds were significantly rescued in the high-dose group (60.83 ± 4.17 dB), the intermediate-high-dose group (62.92 ± 3.72 dB), and the intermediate-dose group (65.42 ± 2.08 dB), while the low-dose group only showed minimal hearing recovery (83.13 ± 5.34 dB) (**Figure 1B**). The ABR thresholds in response to tone-burst stimuli also showed a trend towards better efficacy from low dose to high dose, suggesting a possible dose-dependent effect (**Figure 1B**). In contrast, hearing recovery was not observed in *Otof* ^−/−^ mice receiving AAV1-hOTOF NT or AAV1-hOTOF CT alone (N or C-terminal of human otoferlin) (**Figure S3A**). Further, we detected otoferlin expression in the high-dose group at 1 month after injection. Otoferlin was detected throughout the whole cochlear turns at a rate of 62.29% ± 3.80, 69.62% ± 1.70, and 76.56% ± 3.03 of the IHCs at the apical, middle, and basal turns, respectively (**Figure 1F**-**G**), whereas no otoferlin was detected in the cochlea injected with AAV1-hOTOF NT (**Figure S3B**).

**Figure 1.**
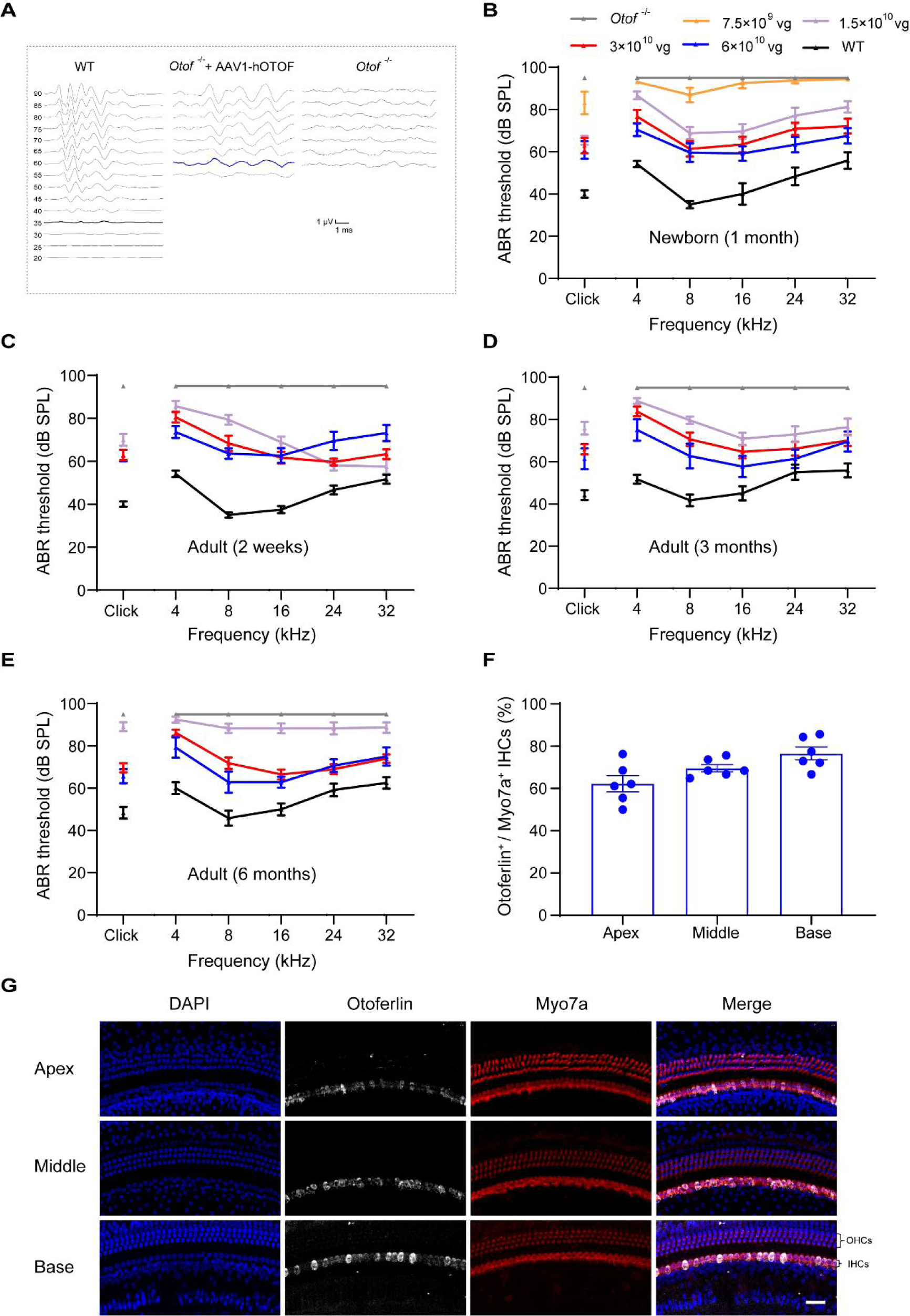
AAV1-hOTOF rescues hearing in *Otof* ^−/−^ mice. **A.** Representative ABR traces in response to broadband click sound stimuli were recorded 4 weeks after therapy injection (6×10^10^ vg/cochlea) at P0-P2. **B.** The ABR thresholds in the injected ear of *Otof* ^−/−^ mice were recorded 1 month after injection at P0-P2. (*Otof* ^−/−^, n = 8; WT, n = 6; 6×10^10^ vg/cochlea, n = 12; 3×10^10^ vg/cochlea, n = 12; 1.5×10^10^ vg/cochlea, n = 12; 7.5×10^9^ vg/cochlea, n = 8). **C.** The ABR thresholds in the injected ear of adult *Otof* ^−/−^ mice were recorded 2 weeks after injection at P30 (*Otof* ^−/−^, n = 8; WT, n = 6; 6×10^10^ vg/cochlea, n = 11; 3×10^10^ vg/cochlea, n = 18; 1.5×10^10^ vg/cochlea, n = 14). **D.** The ABR thresholds in the injected ear of adult *Otof* ^−/−^ mice were recorded 3 months after injection at P30 (*Otof* ^−/−^, n = 8; WT, n = 6; 6×10^10^ vg/cochlea, n = 11; 3×10^10^ vg/cochlea, n = 17, 1.5×10^10^ vg/cochlea, n = 12). **E.** The ABR thresholds of adult *Otof* ^−/−^ mice were recorded 6 months after injection at P30 (*Otof* ^−/−^, n = 8; WT, n = 6; 6×10^10^ vg/cochlea, n = 7; 3×10^10^ vg/cochlea, n = 16, 1.5×10^10^ vg/cochlea, n = 12). **F-G**. Otoferlin expression in the newborn *Otof* ^−/−^ mice injected with 6×10^10^ vg/cochlea observed at 1 month. The percentage of otoferlin-labeled IHCs in the injected ear (n = 6) (**F**). Representative images of IHCs immunostained with otoferlin, including the apical, middle, and basal turns (**G**). Scale bars: 20 μm. Data (**B**-**F**) are displayed as mean ± SEM.

The human inner ear is fully developed in the uterus in contrast to the mouse inner ear which becomes fully mature around 3 weeks postnatally. To evaluate the relevance of our gene therapy approach for humans, we also injected AAV1-hOTOF into adult *Otof* ^−/−^ mice at P30 with different doses via the RWM and then tested their auditory function. At 2 weeks after injection, the click ABR thresholds in injected ear of adult *Otof* ^−/−^ mice were significantly rescued in high-dose group (62.73 ± 2.73 dB), intermediate-high-dose group (63.06 ± 2.40 dB) and intermediate-dose group (70 ± 2.77 dB), respectively (**Figure 1C)**, which was consistent with the hearing improvement in neonatal *Otof* ^−/−^ mice. To assess the duration of hearing recovery, we measured the ABR thresholds in the injected ear of adult *Otof* ^−/−^ mice at 3 months after injection. Compared to the click ABR thresholds at 2 weeks after injection, hearing recovery remained stable in the high-dose group (61.36 ± 4.91 dB) and the intermediate-high-dose group (65.88 ± 2.40 dB), with the best performer recovering to ∼40 dB, which was similar to WT mice. In contrast, in the intermediate-dose group the click ABR thresholds were only moderately increased (75.83 ± 3.07 dB) (**Figure 1D**). In order to observe long-term effects of hearing recovery, we also measured the click ABR thresholds in the injected ear of adult *Otof* ^−/−^ mice at 6 months after injection. Hearing recovery was still observed in the high-dose group (65.71 ± 3.35 dB) and the intermediate-high-dose group (69.69 ± 2.16 dB). However, the click ABR thresholds was 89.17 ± 2.12 dB in the intermediate-dose group (**Figure 1E**). These results support that AAV1-hOTOF RWM delivery partially rescues hearing in adult *Otof* ^−/−^ mice long term with an efficacy that is positively correlated with an increasing dosage.

### Auditory function was not affected in WT mice after AAV1-hOTOF injection

To evaluate whether the injection of AAV1-hOTOF via the RWM has adverse effects on the inner ear and hearing, we examined the auditory function and counted the number of HCs in WT mice. Adult WT mice were randomly assigned to the AAV1-hOTOF-injected or vehicle-injected groups. Limited by the space volume of the cochlea, the maximum dose of AAV1-hOTOF (6×10^10^ vg/cochlea, 2 μL) and an equal volume of vehicle were injected into adult WT mice. The ABR was assessed 2 weeks after injection. Compared with the contralateral uninjected ears, the ABR thresholds showed no changes in the injected ears across all frequencies in the two groups (**Figure 2A**-**B**). Further, we performed immunohistochemistry and counted the number of HCs in order to observe the survival of HCs. There was no statistical difference between the injected ears and contralateral uninjected ears, suggesting that AAV1-hOTOF did not cause obvious adverse effects and was well tolerated in the cochlea (**Figure 2C-F**).

**Figure 2.**
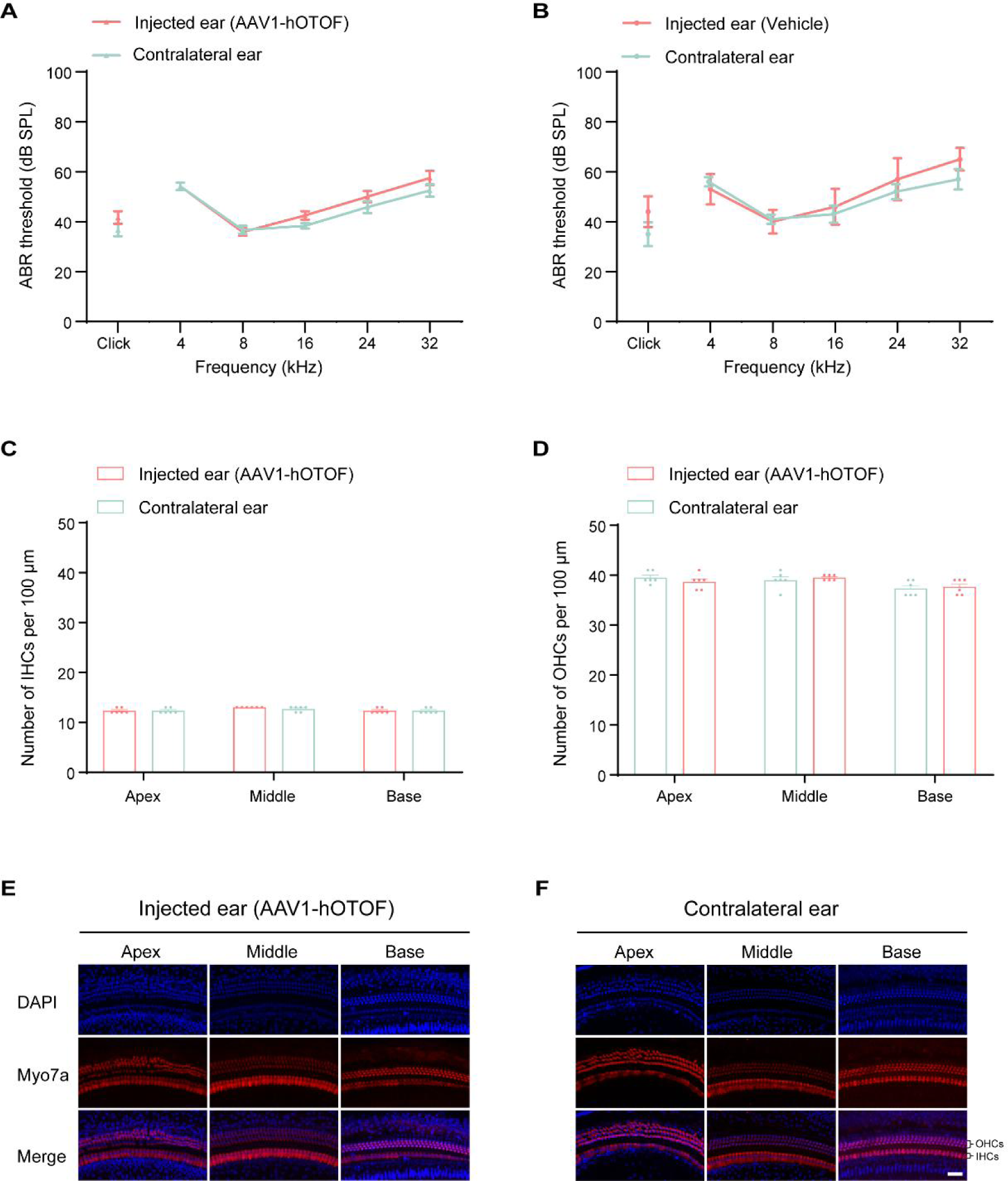
Auditory function was not affected in WT mice after AAV1-hOTOF injection. **A.** The ABR thresholds of AAV1-hOTOF-injected (6×10^10^ vg/cochlea, 2 μL) and contralateral uninjected ears were recorded 2 weeks after injection (n = 6). **B.** The ABR thresholds of vehicle-injected (2 μL) and contralateral uninjected ears were recorded 2 weeks after injection (n = 5). **C**, **D** were quantified for IHCs (**C**) and OHCs (**D**). **E**, **F.** Representative images of Myo7a-labeled HCs from the AAV1-hOTOF-injected (**E**) and contralateral uninjected ears (**F**) were observed at 2 weeks after injection. Scale bar, 20 μm. Data **(A-D**) are displayed as mean ± SEM.

### Biodistribution of AAV1-hOTOF in WT mice after RWM injection

Although local injection of AAV1-hOTOF into the inner ear provided an effective treatment for DFNB9 in mice, few studies have comprehensively reported on the systemic biodistribution of AAV1 in mice following local administration via the RWM. It is necessary therefore to assess the systemic biodistribution of AAV1-hOTOF related to its development in clinical application for gene therapy. To study the biodistribution of AAV1-hOTOF, the total DNA was extracted and analyzed by quantitative polymerase chain reaction (qPCR) targeting AAV1-hOTOF NT to determine the viral genome copy number in the cochlea, brain, and liver as well as other major organs at 6 weeks after injection. As expected, the injected ear had the highest copy number of vector genome (1.2×10^6^ vg/μg DNA) **(Figure S4)**. The viral genome was 10^4^-10^6^ vg/μg DNA in the contralateral uninjected ear, brain, and spinal cord, and at 10^3^-10^4^ vg/μg DNA in the liver, heart, spleen, and blood, whereas in other tissues the vector genome copy number was less than 10^3^ vg/μg DNA (**Figure S4**). In summary, RWM injection yielded a high vector copy number in the cochlea, which was consistent with the treatment efficacy. The detection of a relatively high copy number of viral particles in the central nervous system (CNS) and the liver suggests the necessity of careful safety assessment of potential toxicity in future clinical studies.

### Normal behavior was not affected in WT mice after AAV1-hOTOF injection

To assess whether injected AAV1-hOTOF affects the general condition and behavior of mice, 2 μL of AAV1-hOTOF or vehicle was injected into adult WT mice cochlea, and behavioral tests including the rotarod test, balance beam test, open field test, Y-maze test, and Irwin test were performed.

In the rotarod test there was no significant difference in the falling time or falling speed between AAV1-hOTOF-injected and vehicle-injected groups (*P* > 0.05 for 2 and 4 weeks). Before the injection, the two groups had similar rotarod test baseline results (*P* > 0.05) (**Figure 3A**-**B**). In the balance beam test, all experimental animals succeeded in traversing the balance beam without falling off. Animals’ speed crossing the beam (both 1 cm and 2 cm wide) showed no statistical differences between the AAV1-hOTOF-injected and vehicle-injected groups *(P* > 0.05 for 2 and 4 weeks) (**Figure 3C**-**D**). In the open field test, we measured the distance travelled and time spent in the center region at 4 weeks after injection to quantify their locomotor activity and anxiety-like behaviors. The AAV1-hOTOF-injected mice travelled a similar distance in 30 min as the vehicle-injected group (90.20 ± 6.58 m vs. 98.30 ± 7.87 m, *P* = 0.44) (**Figure 3E)**. The time that AAV1-hOTOF-injected mice spent in the center region (61.62 ± 8.68 s) was comparable to that in the vehicle-injected group (79.65 ± 6.73 s) (*P* = 0.12) (**Figure 3F)**. In the Y-maze test, the number of alterations, number of arm entries, and alteration percentage in the AAV1-hOTOF-injected group were similar to the vehicle-injected group (*P* > 0.05 for all comparisons) (**Figure 3G**-**I**). The Irwin test was performed in mice at 1, 2, 3, and 4 weeks after injection. Between the two groups, no significant behavioral changes were observed in terms of general behaviors, convulsive behaviors and excitability, or reflex capabilities across the duration of the experiment (**Table S1**). No significant changes of the anus temperature were found between two groups after injection (**Figure S5**). Overall, all of the experimental mice were healthy, and no obvious behavioral changes could be attributed to the local injection of AAV1-hOTOF.

**Figure 3.**
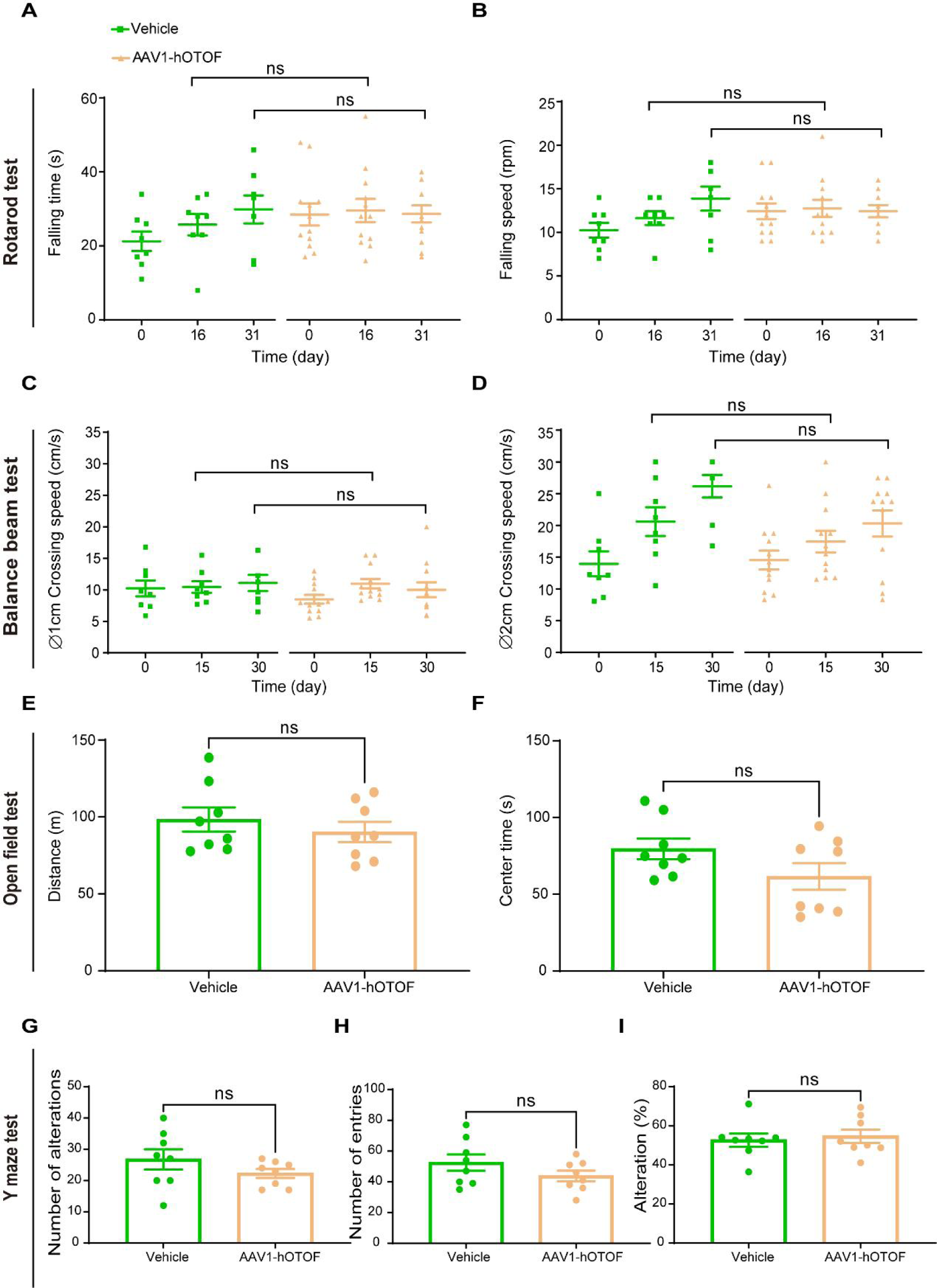
Normal behavior was not affected in WT mice after AAV1-hOTOF injection. **A**, **B.** The rotarod performance of AAV1-hOTOF and vehicle-injected mice in falling time (**A**) and falling speed (**B**). **C**, **D.** The speed of crossing the 1 cm (**C**) and 2 cm (**D**) balance beam were recorded from AAV1-hOTOF and vehicle-injected mice (**A**-**D**, AAV1-hOTOF group, n = 12; vehicle group, n = 8). **E**, **F.** The open field performance of AAV1-hOTOF and vehicle-injected mice in the total travelled distances (**E**) and the time spent in the center region (**F**). **G**-**I.** The Y-maze performance of AAV1-hOTOF and vehicle-injected mice in the number of alterations (**G**), number of entries (**H**), and the percentage of alteration (**I**) (**E**-**I**, AAV1-hOTOF group, n = 8; vehicle group, n = 8). Data (**A**-**I**) are displayed as mean ± SEM. In (**A**-**I**), ns, not significant; \**P* < 0.05; \*\**P* < 0.01; \*\*\**P* < 0.001; \*\*\*\**P* < 0.0001, unpaired *t*-test.

### Abnormal clinical pathology was not observed in WT mice after AAV1-hOTOF injection

To further evaluate the potential systemic toxicity and inflammatory response to AAV1-hOTOF, mice from the AAV1-hOTOF-injected and vehicle-injected groups were phlebotomized at 6 weeks after injection, and the blood was analyzed by routine blood test and serum chemistry. For all routine blood test parameters, there were no statistical differences (*P* > 0.05 for all comparisons) between AAV1-hOTOF-injected group and the vehicle-injected group (**Table S2)**. No evidence of toxicity in serum chemistry was observed, and the values of hepatic function were comparable between the two groups (*P* > 0.05 for all comparisons) (**Table S3**). To evaluate the possible damage in the AAV1-hOTOF-injected and vehicle-injected mice 13 weeks after injection, histologic analysis was performed via hematoxylin and eosin staining in the important organs, including the lung, liver, pancreas, colon, jejunum, kidney, brain, lymph nodes, spleen, bone marrow, skeletal muscle (thigh), and heart. Two vehicle-injected mice showed mild kidney inflammation, and one AAV1-hOTOF-injected mouse showed mild signs of inflammation in the colon. No pathological changes or signs of inflammation or fibrosis were observed in the remaining mice. We conclude that AAV1-hOTOF injection did not cause pathological inflammation (**Figure S6**).

### Delivery of AAV1-GFP via RWM in the non-human primate (NHP) inner ear results in efficient expression in HCs

Vector characterization in the NHP model is an important step prior to clinical trials. We investigated the expression of AAV1-delivered transgenes under the control of the HC-specific promoter *Myo15* in the inner ear of NHP. Two *Macaca fascicularis* were injected (**Table S4**). The first one (animal #1) received AAV1-GFP (2.5×10^11^ vg in 20 μL vehicle) in the left ear via RWM injection using the trans-mastoid approach reported by Andres-Mateos et al.^38^ For the second monkey (animal #2), we injected the left ear with a lower dose (1.5×10^11^ vg in 20 μL vehicle). Four weeks later, the animals were sacrificed and the temporal bone was extracted and decalcified over 2 months.

To evaluate the GFP expression in HCs, 40× images of each mapped frequency region were taken with a confocal microscope (**Figures 4A**, **S7A**-**B**). Significantly, we only detected GFP in the HCs but not in other regions of the cochlea. GFP-positive HCs were counted, and the values were plotted as a percentage of the total number of HCs. In the cochlea with high-dose injection, GFP-positive cells were detected in IHCs (apex, 63.16%; middle, 82.35%; base, 93.75%) and OHCs (apex, 37.82%; middle, 49.58%; base, 27.78%) (**Figure 4B**-**C**). In the cochlea with low-dose injection, GFP-positive cells were also detected in IHCs (apex, 65.38%; middle, 61.54%; base, 87.5%) and OHCs (apex, 37.74%; middle, 61.54%; base 24.24 %) (**Figure 4B**-**C**). In addition, no GFP-positive cells were observed in the contralateral uninjected ear (**Figure S7B**). These results indicate that the AAV1 vector driven by the HC-specific *Myo15* promoter could mediate efficient and specific gene expression in HCs of the NHP inner ear.

**Figure 4.**
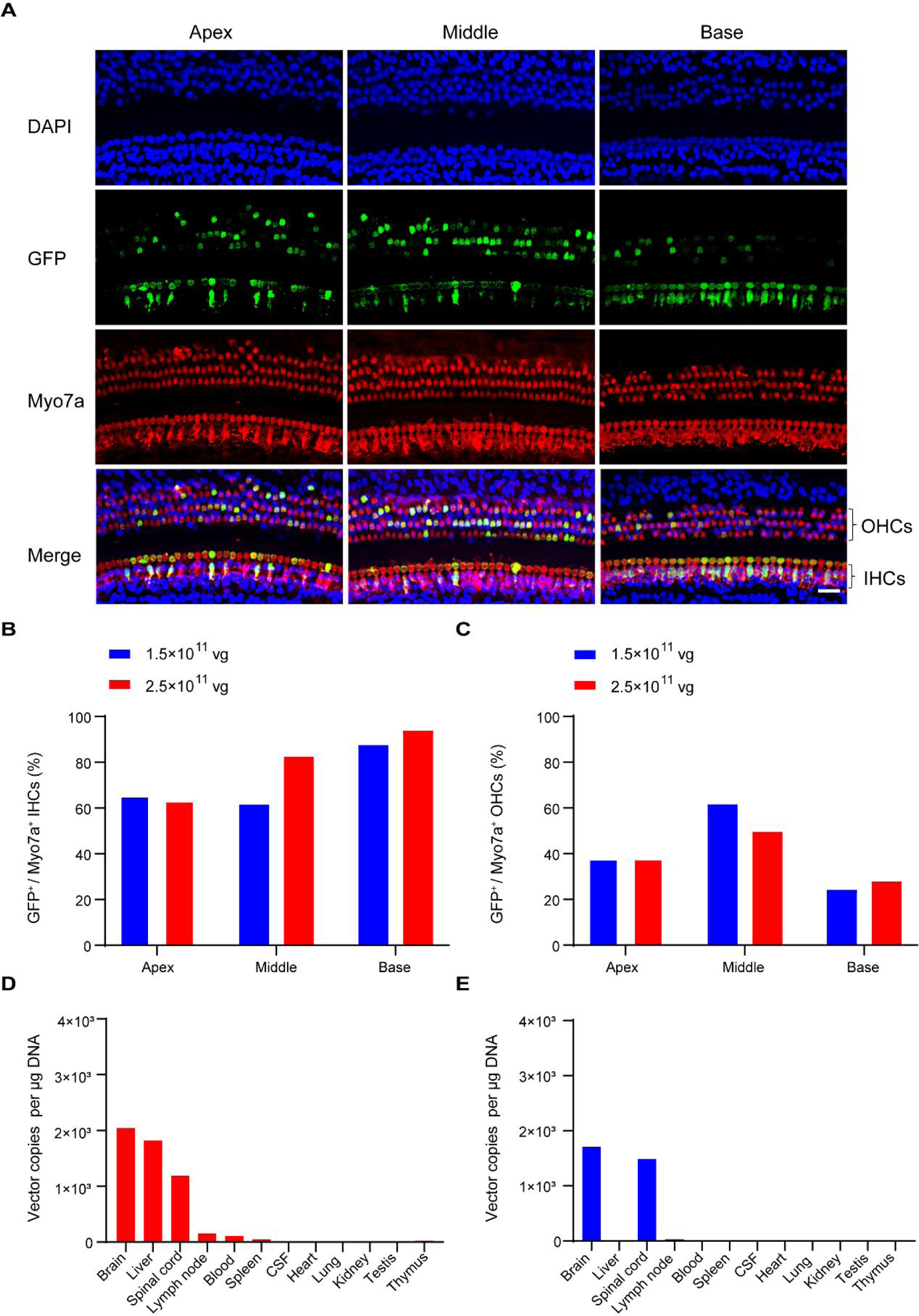
High expression of AAV1-GFP in NHP inner ear hair cells and the vector biodistribution via the RWM route. **A.** Representative images of Myo7a-labeled HCs in the injected ear of NHP at 4 weeks after injection with 2.5×10^11^ vg AAV1-GFP. Scale bar, 20 μm. **B**, **C.** Expression efficiency of GFP in IHCs and OHCs from apex to base in the NHP cochlea injected with 2.5×10^11^ vg and 1.5×10^11^ vg. **D**, **E**. Analysis of AAV1-GFP biodistribution in different organs from NHPs injected with 2.5×10^11^ vg (**D**) and 1.5×10^11^ vg (**E**). Samples were harvested 4 weeks after the injection (n = 2).

### Acute vector tolerability and biodistribution of AAV1-GFP in NHPs

We further evaluated systemic toxicity in NHPs after injection. NHPs were phlebotomized post-surgery (day 14 and day 28), and blood was analyzed for routine blood test (**Table S5**) and serum chemistry (**Table S6**). Systemic toxicity was not detected by serum chemistry or routine blood test in injected NHPs, with the values mostly within the normal reference range based on a previous study,^39^ although a couple of parameters, such as albumin, aspartate aminotransferase, total bilirubin and total protein, demonstrated some an extent of fluctuation.

To study the biodistribution of AAV1-GFP in the injected NPHs, we assessed the viral genome copy number in the blood, brain, liver, and other peripheral tissues at 4 weeks after injection. Compared to other tissues, higher amounts of AAV1 were detected in the brain, spinal, and liver of NHP with a high-dose injection (**Figure 4D**), while AAV1 was minimally detected in the liver of NHP with a low-dose injection (**Figure 4E**). Overall, AAV1-GFP was well tolerated, and we did not observe any apparent systemic acute toxicity related to AAV1-GFP injections.

## Discussion

AAV-mediated gene therapy in humans has been applied successfully to treat genetic diseases, including hemophilia, neurological disorders, and blindness, etc.^40–45^ However, gene therapy for hereditary hearing loss has not reached a clinical stage. Here, in an effort to advance *OTOF* gene therapy towards the clinical stage, we assessed the efficacy and safety of our AAV1-hOTOF agent in animal models. Our data showed that AAV1-hOTOF significantly rescued hearing in *Otof* ^−/−^ mice with profound hearing loss without obvious adverse effects. The hearing rescue was at least 6 months and seemed to positively correlate with an increasing dosage. Our results further demonstrated that the AAV1-transgene driven by the *Myo15* promoter was highly expressed in the HCs of the NHP inner ear and did not cause apparent systemic acute toxicity.

Previous studies have shown that gene replacement is a feasible strategy for gene therapy in *Otof* ^−/−^ mouse models,^15–17^ and this was supported by our data. However, from a safety perspective, the design of our candidate drug was different from previous reports regarding the key constituent elements important in the clinic, especially the AAV delivery vector and promoter. Because there are no cell-specific AAVs for the cochlea currently, AAV1 is commonly applied in clinical trials with good safety records,^29–33^ and HC-specific promoters will be ideal choice for constructing gene therapy drugs for the treatment of DFNB9. For this reason, we selected AAV1 with the tropism primarily in IHCs ^34–36^ as a delivery tool and designed a candidate drug (AAV1-hOTOF) driven by the *Myo15* promoter, to further confine the expression of hOTOF to HCs. We hypothesized that our approach may decrease the ectopic expression of hOTOF via local injection compared to the therapeutic systems used in previous reports.

To test this hypothesis, we injected AAV1-hOTOF into mouse models by the route of RWM. As expected, high expression of otoferlin specifically in HCs was detected, which might be very beneficial for human gene therapy applications by avoiding potential off-target effects. We found that AAV1 was predominately distributed in the inner ears, although AAV1 was also detected in the CNS and liver. One reason for the distribution of AAV1 in other organs may be the cochlear aqueduct, which is a communication channel between the cerebral spinal fluid (CSF) and the perilymph; while another reason may be internal auditory meatus or modiolus, providing probable pathway.^46–50^ In our study, the high-dose injection did not affect the normal behavior, hepatic function, and the normal function other important organs in WT mice, which suggests a profile of safety and tolerability of AAV1-hOTOF for future use in clinical studies.

NHPs are phylogenetically close to humans in physiology, anatomy, immunology, and neurology, all of which make them excellent experimental models to assess the toxicity and pharmacology of vector systems prior for consideration for preclinical studies.^27, 38, 51, 52^ In our study, AAV1-GFP showed good efficiency in transducing the cochlear HCs in NHPs. Compared to the previous study,^38^ we detected relatively higher GFP expression in HCs with no transduction of other cell types along the length of the cochlea following RWM injection in NHPs. There are several possible explanations for the expression discrepancy between our study and the previous study. First, this could be due to pre-existing immunity to AAV in the perilymph that affects the transduction of the cochlea because the significant correlation between the titers of antibody and the levels of GFP expression has been shown previously.^53^ It might be necessary to perform the detection of neutralizing antibodies against AAV before injection of AAV-mediated gene therapy system. Second, the small anatomical size and limited exposure of the oval and round window may result in difference in AAV delivery during the surgical procedure. Third, other factors that may explain the discrepancies include the promoter, vector quality, volume, duration of study, the animal age at the time of surgery, and the animal species.^54^

Biodistribution in the CNS and systemic biodistribution were also quantified in our study. In a previous study, a patent cochlear aqueduct together with local AAV injection resulted in the transduction of cerebellar Purkinje cells in mice.^47^ While there is still a debate about the patency of the cochlear aqueduct in NHPs and humans,^55, 56^ the majority of adult human temporal bones are more likely to be occluded with loose connective tissue.^57^ However, some studies thought that the cochlear aqueducts are patent in humans, which does not preclude the possibility of the viral vectors reaching the CNS after inner injection.^58, 59^ In our study, no GFP expression was detected in the contralateral uninjected ear, and AAV1 genome copies were detected at a low level in the CNS tissue, followed by the spinal cord and the liver, while other peripheral tissues had relatively lower vector copies in NHPs. There was no evidence of toxicity observed by serum chemistry and routine blood tests, although a few differences were detected in some parameters, suggesting the safety of the AAV1 vector carrying the *Myo15* promoter and the *GFP* or *OTOF* transgene, which may further strengthen the safety profile. We will continue to expand the number of NHPs with longer follow-up in future studies.

The human cochlea is fully developed in utero, while the rodent cochlea is not mature until 3 weeks after birth.^60^ The cochlea is a relatively isolated organ compared to systemic injection, and local injection in the inner ear tends to result in a high drug concentration inside the cochlea and to minimize the distribution in other organs. Therefore, successful gene therapy in the mature inner ear is more relevant for translational application in the clinic. The RWM route, delivering therapeutic agents into the scala tympani, is one of the most common approaches for gene therapy in adult mice,^61–63^ but a novel delivery strategy that combines RWM injection with canalostomy has demonstrated highly-efficient transduction throughout the cochlea and minimal hearing impairment in rodents.^61, 62^ In previous studies, the invasive trans-mastoid approach of RWM injection after making a fenestra in the oval window was implemented successfully in NHPs.^38, 64^ The human cochlea has more space to operate than the NHP cochlea. We thus expect the approach of RWM injection with canalostomy or making a fenestra in the oval window to be a feasible gene transfer method for gene therapy to treat deafness in humans.

In summary, our study established a therapeutic agent – AAV1-hOTOF – for DFNB9 and demonstrated its efficacy and safety in mouse models. Additionally, we explored the transduction and tolerance of AAV1-GFP in NHPs. These findings strongly support the clinical development of AAV1-hOTOF.

## Materials and Methods

### Mice

Details of the *Otof* ^−/−^ mice, established by BIOCYTOGEN (Beijing, China), are described in our previous study.^15^ WT mice (129 strain) were used in the experiments to evaluate the AAV1-hOTOF. All mice were raised in the Department of Laboratory Animal Science of Fudan University, with free access to food and water and 12-h light/dark cycles. All animal experiments were performed according to guidelines regarding institutional animal welfare and were approved by the Animal Care and Use Committee of Fudan University, China.

### Construction of dual-AAV strategies

For the therapeutic agent constructs, the full-length human *OTOF* CDS (NM_001287489.2) was split at the exon 21-exon 22 junction site, which was divided into 5′-terminal and 3′-terminal segments referring to previous study.^17^ The human *OTOF* CDS was synthesized by Sangon Biotech Co., Ltd. (Shanghai, China). The 5’-terminal and 3’-terminal segments were packaged into the AAV1 capsid. Both otoferlin dual-5’ AAV1-TS and 5’ AAV1-Hyb half-vectors carried the CAG or *Myo15* promoter, the 5’-terminal segment of the *OTOF* CDS, and a splice donor (SD) sequence. ^37^ The dual-3’ AAV1-TS and 3’ AAV1-Hyb half-vectors carried a splice acceptor (SA) sequence, the 3′-terminal segment of the *OTOF* CDS, a woodchuck hepatitis virus post-transcriptional regulatory element (WPRE), and a bovine growth hormone polyadenylation sequence (pA). In addition, the highly recombinogenic sequence derived from the F1 phage (AK) or alkaline phosphatase (AP) ^37^ was added downstream of the SD sequence and upstream of the SA sequence in the dual-5’ AAV1-Hyb and 3’ AAV1-Hyb half-vectors. The therapeutic system of the dual-AAV1-Hyb (AK) approach in which the human *OTOF* CDS was driven by *Myo15* promoter was called AAV1-hOTOF for short. The N and C-terminal of human *OTOF* packaged by AAV1 were called AAV1-hOTOF NT and AAV1-hOTOF CT, respectively.

### Virus production

The AAV2 inverted terminal repeats (ITRs) were used as a packaging signal for packaging with the AAV1 serotype. The AAVs used in this study include AAV1-CAG-hOTOF NT-AK, AAV1-CAG-hOTOF NT-AP, AAV1-CAG-hOTOF NT-TS, AAV1-Myo15-hOTOF NT-AK, AAV1-Myo15-hOTOF NT-AP, AAV1-Myo15-hOTOF NT-TS, AAV1-hOTOF CT-AK, AAV1-hOTOF CT-AP, AAV1-hOTOF CT-TS and AAV1-Myo15-GFP-WPRE-bGH-polyA. The vector plasmid together with capsid and helper plasmids were transiently transfected into HEK293T cells to produce viral particles. The viruses were commissioned from PackGene Biotech (Guangzhou, China). The virus genome titer was determined by qPCR, and they were stored at −80°C and thawed just before injection.

### Expression and recombination of otoferlin in HEK293T cells

The HKE293T cells were cultured in high-glucose DMEM with 10% FBS and Penicillin/Streptomycin (Thermo Fisher Scientific, #11965092) in a humidifed incubator free from mycoplasma and chlamydia at 5% CO2 and 37°C. Based on three different recombination strategies, AAV1-CAG-hOTOF NT and AAV1-hOTOF CT were used to infect HEK293T cells by MOI = 2×10^5^. After 48 h, the cells were lysed in cold RIPA bufer (Beyotime, #P0013B) containing PMSF (Beyotime, #ST506,) to release otoferlin, and the total protein concentration was determined using a BCA protein assay kit (Beyotime, #P0012S). The protein samples were separated by SDS-PAGE (4-12% Bis-Tris gels, Genscript, #M41210C) with MOPS buffer (Tris-MOPS-SDS Running Buffer, Genscript, #M00138) and were transferred onto polyvinylidene fuoride membranes. The membranes were probed with the primary antibodies rabbit IgG anti-OTOF (1:2000 dilution, Thermo Fisher Scientific, #PA5-52935), rabbit IgG anti-GAPDH (1:1000 dilution, CST, #5188), and followed by incubation with the anti-rabbit IgG-HRP secondary antibody (1:2000 dilution, Beyotime, #A0239). Finally, the secondary antibodies were detected using Western Blot ECL Blotting Substrate (Thermo Fisher Scientific, #34580) with an Tanon 4600 system.

### Inner ear injection

The surgical procedure of inner ear injection via the RWM was as described previously.^15, 65^ Briefly, P0-P2 mice were anesthetized by hypothermia on ice. Cochleostomy was performed by post-auricular incision to expose the cochlear bulla, and a glass micropipette (WPI, Sarasota, FL) connected to a Nanoliter Microinjection System (WPI) was used to manually deliver the AAV1-hOTOF into the inner ear via RWM. The operation was conducted only in the right ear of each animal. The incision was closed with sutures after the injection, and the mice were placed on a heating pad (37℃) for resuscitation and then returned to their cage.

Adult mice were anesthetized with ketamine (100 mg/kg) and xylazine (10 mg/kg) via intraperitoneal injection. The operated post-auricular region was exposed by shaving and was disinfected with 10% povidone-iodine. Surgery was performed under an operating microscope. We made a 10-mm post-auricular incision to access the temporal bone and subsequently drilled a small hole in the tympanic bulla to expose the round window. A glass micropipette held by a Nanoliter Microinjection System was used to penetrate the RWM and left it open for a few minutes until no obvious perilymph leakage was observed. We performed the vector or vehicle (PBS with 0.001% PF68) delivery with an injection rate of 5 nL/s controlled by Nanoliter Microinjection Controller (WPI). The RWM niche was sealed with a small plug of muscle quickly after the injection, then the otic bulla was sealed with muscle using tissue adhesive. The skin was closed with sutures.

### Hearing test

The ABR was recorded in a soundproof chamber using the TDT BioSigRP system (Tucker-Davis Technologies, Alachua, FL, USA) as previously described.^14, 15, 66^ Mice of either sex were anesthetized through intraperitoneal injection with xylazine (10 mg/kg) and ketamine (100 mg/kg).^15, 66^ The recording electrode, the reference electrode and the ground electrode were inserted at the mastoid portion, the subcutaneous tissues of the vertex, and the rump of the mice, respectively. We examined the mice to exclude otitis media or abnormal formation of cerumen in the ear canal before recording. The sound level was decreased in 5 dB steps from 90 dB to 20 dB sound pressure level (SPL). Tone burst acoustic stimuli were presented at 4, 8, 16, 24, and 32 kHz. The ABR responses were amplified (10000 ×), filtered (0.3-3 kHz), and averaged (1024 responses) at each SPL. The ABR threshold was defined as the lowest SPL level at which any wave could be visually detected and repeated. The acoustic stimuli thresholds were defined by two independent observers.

## Behavioral test

### Rotarod test

A mouse rotary rod instrument was used to assess the ability of motor coordination and motor learning of the mice.^67^ In the training session, the mice were placed on a rotating rod with a diameter of 3 cm to acclimate to the equipment, and the speed was set to 8 rpm. In the test session, mice were placed on a slowly moving rod and the rotation speed was accelerated from 4 rpm to 40 rpm in 5 min. The time and the rotation speed when the mouse fell off were recorded. The apparatus was cleaned to prevent the residual information from the previous mouse from affecting the test results.

### Balance beam test

The balance beam test was used to assess the motor coordination and balance of mice.^67^ The beam (1 or 2 cm wide, 100 cm length) was elevated 50 cm. A black plastic box (30 cm × 15 cm × 15 cm) was set as a nest at the end of the beam for motivating the animal to cross the beam. Cloth was placed under the equipment to protect the animals from injury if they fell off. The balance beam was cleaned with 75% alcohol and then water between trials. Before the test, the mouse was allowed to rest in the box for 3 min, then placed on the beam 60 cm from the endpoint. After the test, the time and the average movement speed of mice to cross the beam were recorded. If the animal fell off, the distance and time that the animal actually crawled on the beam were recorded, then the mouse was placed in the box to rest for 1 min, and the previous steps were repeated three times.

### Open field test

The open field test was used to assess locomotor activity and anxiety-related behaviours of mice.^68, 69^ During the test, each mouse was placed in the center of a plastic chamber (50 cm × 50 cm × 40 cm), and the movement was recorded for 30 min with a video tracking system. After the test, the mouse travelled distance and the time in center region were analyzed. The chamber was cleaned with 75% alcohol and then water to avoid olfactory cues.

### Y-maze test

The Y-maze consisted of three equiangular arms that were labelled A, B, and C and was used to evaluate the spatial memory of mice.^70^ Mice were placed in the center of the maze and allowed to visit three arms freely for 8 min. The number of arm entries and alternations were recorded. The alteration was determined from successive consecutive entries to the three different arms, such as ABC, BCA, BAC, etc. The alteration was expressed as the percentage of alternations and was calculated as (alternations/ [total arm entries – 2])×100.

### Irwin test

The Irwin test was developed to detect any deleterious effects of a new drug on general behavior. Each mouse was observed in a transparent box, and the observation consists of the general behavior (spontaneous exploration, grooming, smelling its congeners, normal resting state, alertness, distending/oedema, bad condition, moribund, and dead), convulsive behavior and excitability (spontaneous activity, restlessness, fighting, writhing, tremor, stereotypy, twitches / jerks, Straub, opisthotonos, and convulsion), and reflex capabilities (startle response, touch reactivity, vocalization, abnormal gait, corneal reflex, pinna reflex, catalepsy, grip reflex, pulling reflex, righting reflex, body tone, and pain response). The incidence rate and severity of symptoms were analyzed. The rectal temperature of each mouse was measured after observation. The behavioral tests including the rotarod test, balance beam test, open field test, Y-maze test, and Irwin test were performed by WuXi AppTec Co., Ltd. (Shanghai, China) and PharmaLegacy Laboratories (Shanghai, China).

## Immunohistochemistry

Cochleae were harvested immediately after the mice were sacrificed and they fixed in 4% paraformaldehyde (PFA, Sangon Biotech, #E672002-0500,) at 4°C overnight. A small hole was then made at the apex of the cochlea to facilitate PFA irrigation, and the cochleae were decalcified in 10% EthyleneDiamine TetraaceticAcid (EDTA, Sangon Biotech, #E671001-0500) for 2-3 days. The basilar membrane from the decalcified cochlea was dissected and cut into pieces for immunofluorescence. These membranes were infiltrated with 1% Triton X-100 (Sangon Biotech, #A110694) in PBS, and blocked in 5% donkey serum (Sigma-Aldrich, #D9663) for 1h at room temperature. After that, the tissues were incubated with mouse IgG1 anti-otoferlin (1:200 dilution, Abcam, #ab53233) and the rabbit anti-Myo7a (1:500 dilution, Proteus BioSciences, #25-6790) primary antibodies overnight at 4℃, followed by three washings with PBS. The appropriate Alexa Fluor 555-conjugated anti-mouse IgG1 (1:500 dilution, Invitrogen, #A21127) and Alexa Fluor 647-conjugated anti-rabbit (1:500 dilution, Invitrogen, #A31573) secondary antibodies were then incubated for 2 h at room temperature. The cell nuclei were stained with DAPI (Sigma-Aldrich, #F6057). Images were collected on a Leica TCS SP8 laser scanning confocal microscope with a 40× objective. The images were processed and analyzed with ImageJ software. For IHC and OHC counting, the numbers of Myo7a^+^ HCs and the numbers of GFP^+^/Myo7a^+^ cells were counted in every non-overlapping 100 µm region of the apical, middle, and basal turns of the basilar membrane.

## Tissue biodistribution of vector DNA

Mice were anesthetized with 5% isoflurane and immediately sacrificed by decapitation. Different tissues and organs, including the lymph nodes, pancreas, spinal marrow, thymus, heart, liver, spleen, lung, kidney, testicle, and brain, were carefully dissected and separated. Viral DNA was extracted and purified from tissues using a QIAamp DNA Mini Kit (Qiagen, #51304) following the manufacturer’s instructions and were dissolved in nuclease-free water. The DNA concentration was measured by spectrophotometric analysis using a NanoDrop spectrophotometer (Thermo Fisher Scientific, Waltham, MA, USA). A sufficient volume was calculated such that 500 ng DNA was used for each PCR reaction. Real-time qPCR (RT-qPCR) was used to determine the copy number of AAV1-hOTOF NT in different tissues on a QuantStudio 1 Real-Time PCR instrument (Thermo Fisher Scientific, #INS-0049) using Taqman probes and primers for the 5′-terminal (1-2523 bp) segment of OTOF and AK region and the AceQ qPCR Probe Master Mix (Vazyme, #Q112-02). The forward primer and reverse primer were used for these experiments (*Myo15*-Otof-AK-N forward: 5’-CTGAGGCTGTGCCAGAACT-3’; *Myo15*-Otof-AK-N reverse: 5’-GCAAAATCCCAGAAACGCAAGAG-3’; *Myo15*-SD-AK probe: 5’FAM-TCCTGGCGGACGAGGTAAGTATCAAGG-3’BHQ1). Each 20 μL reaction mixture contained 12.5 μL of AceQ qPCR Probe Master Mix, 0.5 μL of Rox dye (type II), 5.75 μL of RT-PCR grade H_2_O, 0.5 μL (10 μM) each of forward and reverse primer, 0.25 μL (10 μM) of probe primer, and 5 μL (100 ng) of DNA sample. All samples were run in triplicate. Two non-template control reactions (20 μL) and one RNase/DNase-free H_2_O blank (20 μL) were included in each run. The 96-well plates were sealed with an optically clear seal (Applied Biosystems), and the PCR cycling conditions included 5 min at 95°C, and then 40 cycles of denaturation for 15 s at 95°C and annealing and extension for 30 s at 60°C. The concentrations of PCR product were interpolated from the CT values, and triplicate concentration values were averaged.

## Routine blood test and serum chemistry in mice

Mice were anesthetized with 5% isoflurane and immediately sacrificed. Blood samples (1.5-2.0 mL) were taken from mice, which were used for routine blood test and serum chemistry. Routine blood test and serum chemistry were performed by WuXi AppTec Co., Ltd. (Shanghai, China).

## Hematoxylin-eosin staining

Samples from mice tissues and organs were fixed in 10% formalin buffer, embedded with paraffin, sectioned, stained with hematoxylin and eosin, and evaluated by experienced veterinary anatomic pathologists. Hematoxylin-eosin staining was performed according to the standard protocols at PharmaLegacy Laboratories (Shanghai, China).

## NHPs

*Macaca fascicularis* were kindly provided and accommodated by PharmaLegacy Laboratories (Shanghai, China). The NHP experiments were approved by the Institutional Animal Care and Use Committee of Fudan University and the Shanghai Medical Experimental Animal Administrative Committee. The subjects used for experimentation were 5∼7-year-old males. Every animal was healthy and without ear disease or otologic surgery history, such as ear infections, signs of imbalance, or other risk factors for the loss of inner or middle ear function. A psychologically and physically comfortable environment was provided for the animals with a room temperature of 18-28 °C, humidity of 30-70%, and a 12 h light/dark cycle.

## Venous blood and CSF sampling in NHPs

Blood was collected intravenously at 14– and 28-days post-surgery after sterilizing the epidermis of the femoral vein or saphenous vein with 75% alcohol in sedated animals. A standard vacutainer needle was inserted into the vein, and a swab was used to apply pressure to the vessel after the removal of the needle to achieve hemostasis. NHP serum chemistry and routine blood test in the monkeys were performed by Shanghai PharmaLegacy Laboratories (Shanghai, China).

CSF was obtained from the cisterna magna prior to the sacrifice of the monkeys. Atropine was given prior to the operation. Once general anesthesia was established, we removed the hair from the epidermis of the cisterna magna and disinfected the skin with 75% alcohol. After that, the neck of the animal was bent to better expose the area. A 23G needle was used to collect CSF sample.

## RWM injection surgery in NHPs

The protocol was performed according to a previous study.^38^ After the general anesthesia was established, the animal was intubated with a reinforced endotracheal tube. The surgical region was exposed by shaving and then disinfected with 10% povidone-iodine. A 25 mm semilunar incision was made, and the mastoid cortex was exposed. Bipolar electrocautery was used for meticulous hemostasis during the operation. Self-holding retractors was placed, and the surgical microscope was used. The external auditory canal wall, tympanic membrane, the facial nerve, and the chorda tympani were protected during the surgery. After the round window niche and the stapes footplate were visualized, a fenestration in the oval window was made with a needle and observed for a few minutes until no obvious perilymph leakage was found. The bony overhangs of the round window niche were gently removed to expose the RWM.

A Nanoliter Microinjection System (WPI) connected to an injection needle was used to deliver the AAV1-GFP at a rate of 1 μL/min for about 20 min. A total of 20 μL AAV1-GFP was microinjected into the scala tympani through the RWM. After the injection, the RWM and the fenestra in the stapes footplate sealed with muscle. The mastoid cavity was blocked with bone meal and adjacent autologous tissue, and the skin was closed with sutures. During the postoperative care period, animals were carefully monitored and in good care.

## Immunohistochemistry and histology experiments of NHPs

The protocol was performed according to a previous study ^38^. All animals were euthanized without recovery from the deep anesthesia. In brief, the carotid artery was exposed and adequately perfused with saline, followed by formalin solution to achieve full irrigation and preliminarily fixation of the inner ear sample. Cochlear samples and samples from different organs were harvested and processed for histological and/or biodistribution studies.

The cochlear samples were fixed in 4% paraformaldehyde (PFA, Sangon Biotech, #E672002-0500,) and decalcified in 10% EDTA (EDTA, Sangon Biotech, #E671001-0500) at 4°C with regular trimming for about 2 months. The basilar membrane collected from the cochlea was permeabilized with 1% Triton X-100 (Sangon Biotech, #A110694) in PBS and blocked in 5% donkey serum (Sigma-Aldrich, #D9663) for 1h at room temperature. DAPI (Sigma-Aldrich, #F6057) was used to stain the cell nucleus. The primary antibodies were rabbit anti-Myo7a (1:500 dilution, Proteus BioSciences, #25-6790) and chicken IgY anti-GFP (ab13970, 1:500 dilution, Abcam), and the following secondary antibodies were Alexa-conjugated donkey anti-rabbit IgG [H + L], Cy^TM^3 (1:500 dilution, Alex Jackson ImmunoResearch, #711-165-152) and Alexa Fluor 488-conjugated anti-chicken IgY (A11039, 1:500 dilution, Invitrogen). A Leica TCS SP8 laser scanning confocal microscope was used to collect the fluorescent z-stack images (40× objective). The maximum intensity projections of the optical confocal sections are shown in the figures. The images were processed and analyzed using the ImageJ software. For the quantification of GFP-positive cells, all images obtained with the 40× objective of every mapped frequency region (non-overlapping) were manually quantified along the full length of the basilar membrane by two independent researchers who were blinded to the treatment. The percentages were obtained by dividing the number of GFP-positive cells by the total number of Myo7a positive HCs (IHCs and OHCs were calculated separately).

## Tissue DNA extraction and qPCR

Different organ tissues were harvested, weighed and homogenized in buffered Proteinase K and incubated overnight at room temperature, followed by centrifuging for 10 min at 10,000 × *g* at 4°C to a final concentration of 100 µg/ml. Total DNA was extracted using a DNA mini-kit (Qiagen, #51304) according to the manufacturer’s instructions. RT-qPCR was carried out using the QuantStudio 1 Real-Time PCR Detection System (Thermo Fisher Scientific, #INS-0049). Forward primer: 5’-GCTATTGCTTCCCGTATGGC-3’; Reverse primer: 5’-GGAAAGGAGCTGACAGGTGG-3’. Each PCR reaction contained 12.5 μL of SYBR Premix Ex TaqⅡMix, 8.5 μL of ddH2O, 1 μL (10 μM) each of the forward and reverse primers, and 2 μL of the DNA sample. All samples were run in triplicate. Two non-template control reactions (25 μL) and one RNase/DNase-free H_2_O blank (25 μL) were included in each run. The 96-well plates were sealed with an optically clear seal (Applied Biosystems), and the reactions were carried out with an initial incubation at 95°C for 30 s, followed by 40 cycles of denaturation at 95°C for 5 s, annealing at 60°C for 30 s, and extension at 60°C for 30 s. All reactions were carried out in triplicate, and the ‘non-template’ controls containing water instead of template DNA, were included in every PCR run.

## Statistical analyses

Data are depicted as mean and standard error of the mean (SEM). Statistical analysis was performed and plotted using GraphPad Prism 9 (GraphPad Software). An unpaired t-test was used to test for statistical significance between two non-paired normally distributed data groups. Moreover, the two-way ANOVA test followed by Tukey’s multiple comparisons test was used to test for statistical significance in a randomized block design. Statistical significance is indicated in the figures: ns., not significant; \**P* < 0.05; \*\**P* < 0.01; \*\*\**P* < 0.001; \*\*\*\**P* < 0.0001.

## DATA AVAILABILITY

All study data generated in the article and/or supplemental information during the current study are available from the corresponding author on reasonable request.

## ACKNOWLEDGMENTS

This work was supported by the National Natural Science Foundation of China (82225014, 82171148, 82192864), the National Key R&D Program of China (2020YFA0908201, 2021YFA1101302), Shanghai Municipal Health Commission (20224Z0003), Science and Technology Commission of Shanghai Municipality (21S11905100), Shanghai Municipal Education Commission (2023ZKZD12), the Shuguang Program (20SG08) supported by Shanghai Municipal Education Commission and Shanghai Education Development Foundation, and Fudan University (yg2022-23). We also acknowledge the Ines-Fredrick Yeatts Inner Ear Research Fund (Z.Y.C).

## AUTHOR CONTRIBUTIONS

Y.S., H.L., and B.C. developed and supervised the project. L.Z., H.W., M.X., J.W., and J.L., performed the experiments, L.Z., H.W., M.X., and H.T. analyzed the data for the project and wrote the manuscript. Y.S., Z.C., L.Z., Y.C., D.W., Z.G., J.W., H.T., S.H., B.Z., reviewed and revised the manuscript. All authors read and approved the final manuscript.

## DECLARATION OF INTERESTS

The authors have no conflicts of interest to declare.

## Notes

### Competing Interest Statement

The authors have declared no competing interest.

