## Supplemental Information for "Preclinical evaluation of the efficacy and safety of AAV1-hOTOF in mice and non-human primates"

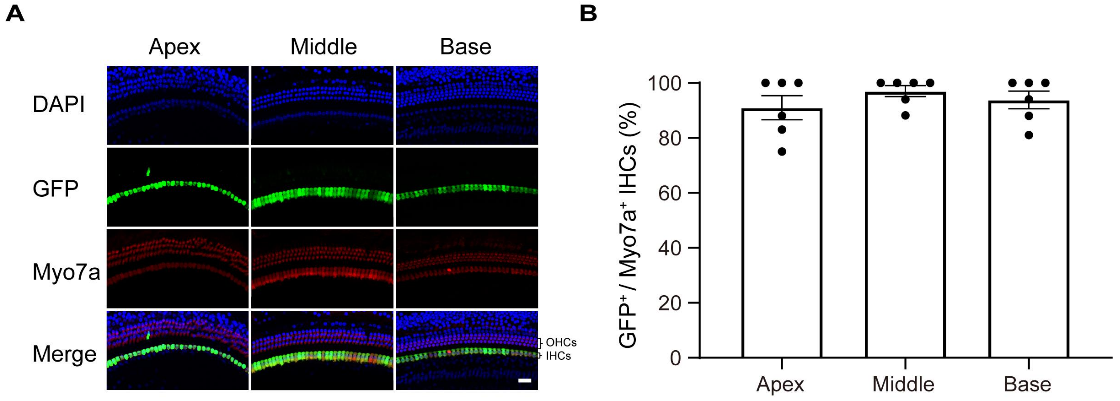


**Figure S1. Representative images of the expression of AAV1-GFP in inner hair cells (IHCs) in WT mice. A**. Representative images of IHCs immunostained with GFP in the injected ears, including the apical, middle, and basal turns. **B**. Percentage of GFP-positive IHCs in the injected ears (n = 6). IHCs were labeled by Myo7a. Data are displayed as mean ± SEM.

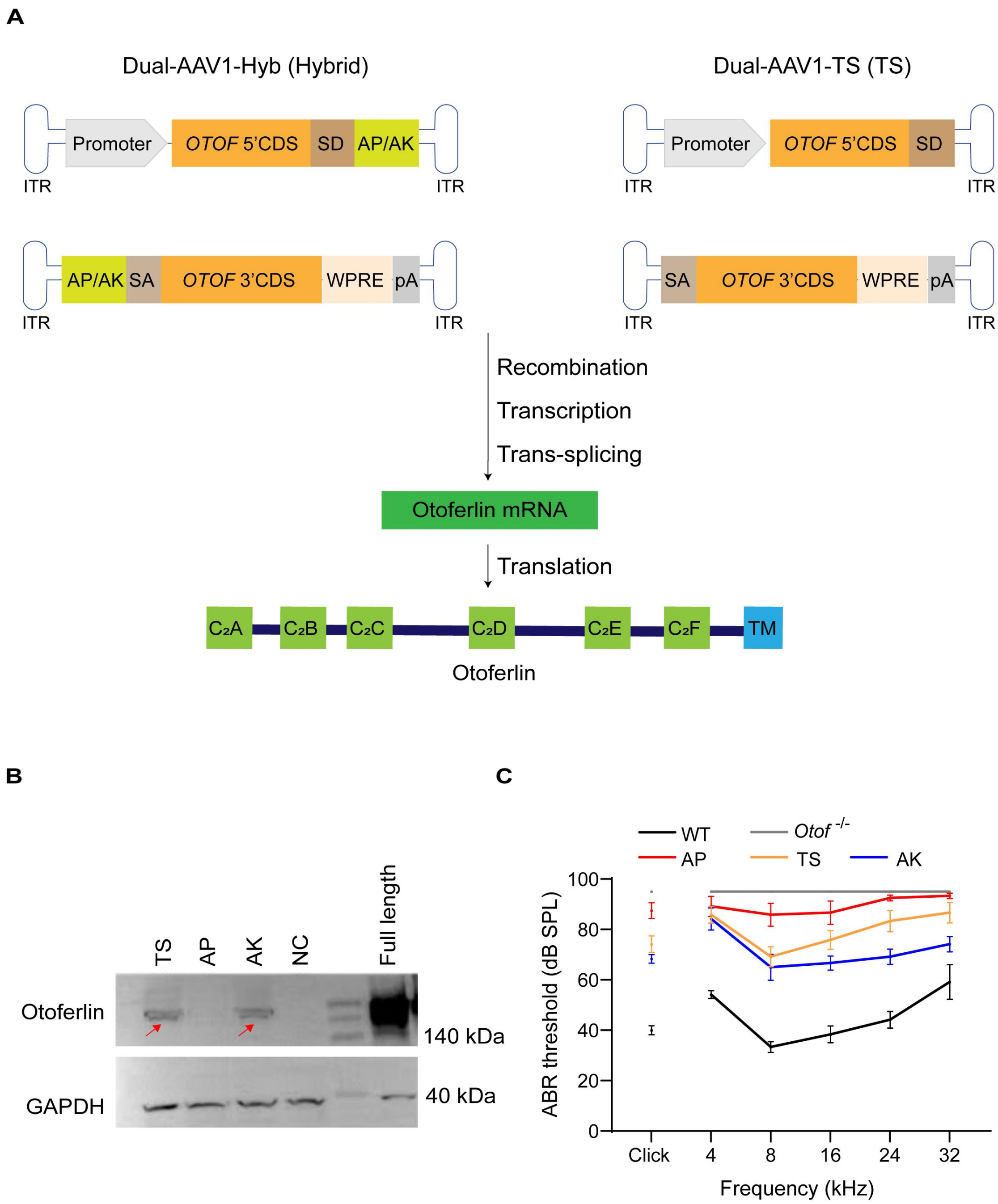


**Figure S2. Identification of human otoferlin recombination. A**. Schematic representation of nucleic acid-based strategies for the recovery of full-length human otoferlin. Reference Transcript ID: NM_001287489.2. In the dual-AAV1-TS strategy, the recombination is mediated by inter-molecular or intra-molecular recombination of inverted terminal repeats (ITRs). In the dual-AAV1-Hyb (AP) and dual-AAV1-Hyb (AK) strategy, the recombination is mediated by ITRs and/or homologous recombination of the recombinogenic AP or AK sequence. SD (splicing donor) and SA (splicing acceptor) sites facilitate the splicing of *OTOF* via trans-splicing. **B.** The expression of full-length human otoferlin protein *in vitro*. The expression of recombinant protein mediated by different strategies (AP, AK, and TS) was verified by western blot. The red arrow indicates otoferlin. The full-length otoferlin was represented by "Full length", while "NC" indicated the negative control in which the cells were not transfected. **C.** The auditory restoration due to different strategies in *Otof* $^{-/-}$

mice (AP, AK, and TS; the dose was $2\times10^{10}$ vg). Abbreviations: CDS, coding sequence; pA, poly-adenylation signal; SD, splicing donor; SA, splicing acceptor; AP, alkaline phosphatase recombinogenic region; AK, F1 phage recombinogenic region; WPRE, woodchuck hepatitis virus post-transcriptional regulatory element.

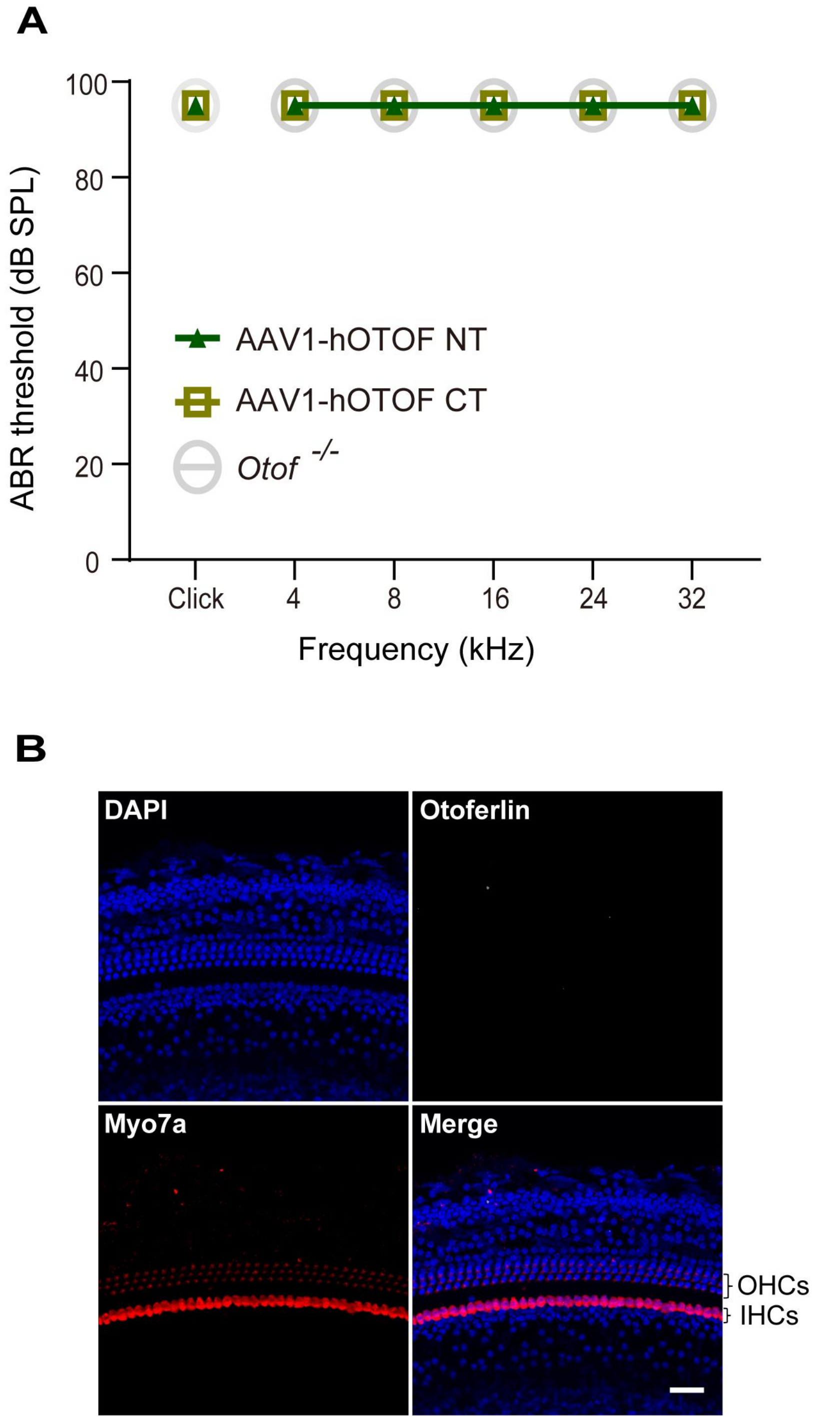


**Figure S3. Auditory function and otoferlin expression in *Otof* $^{-/-}$ mice injected with AAV1-hOTOF NT or AAV1-hOTOF CT alone. A.** ABR thresholds of mice injected with AAV1-hOTOF NT or AAV1-hOTOF CT alone (AAV1-hOTOF NT, n = 5; AAV1-hOTOF CT, n = 5; *Otof* $^{-/-}$, n = 8). **B.** Otoferlin expression in the injected ear of *Otof* $^{-/-}$ mice injected with AAV1-hOTOF NT alone. Scale bar, 20 μm.

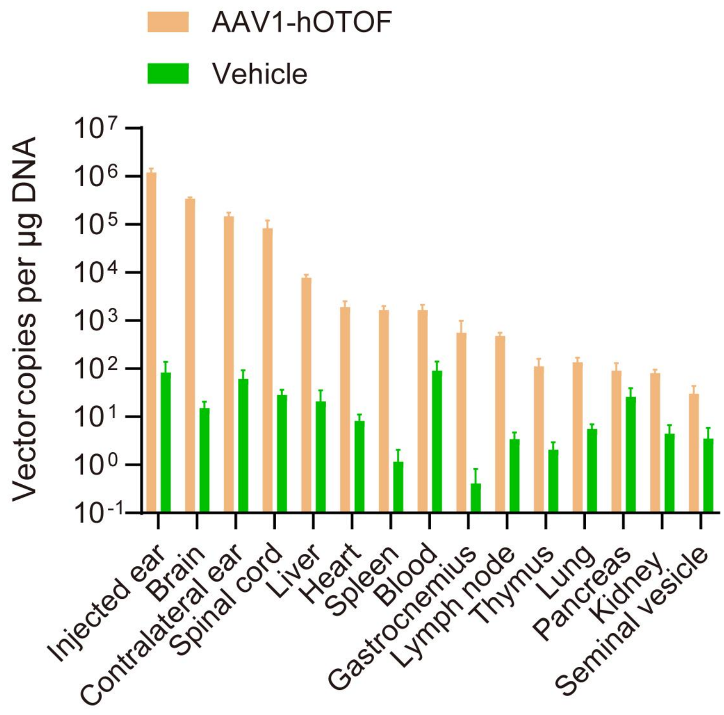


**Figure S4. Analysis of AAV1-hOTOF biodistribution in different organs in WT mice.** Samples were harvested 6 weeks after injection (AAV1-hOTOF group, n = 12; vehicle group, n = 8). Data are displayed as mean ± SEM.

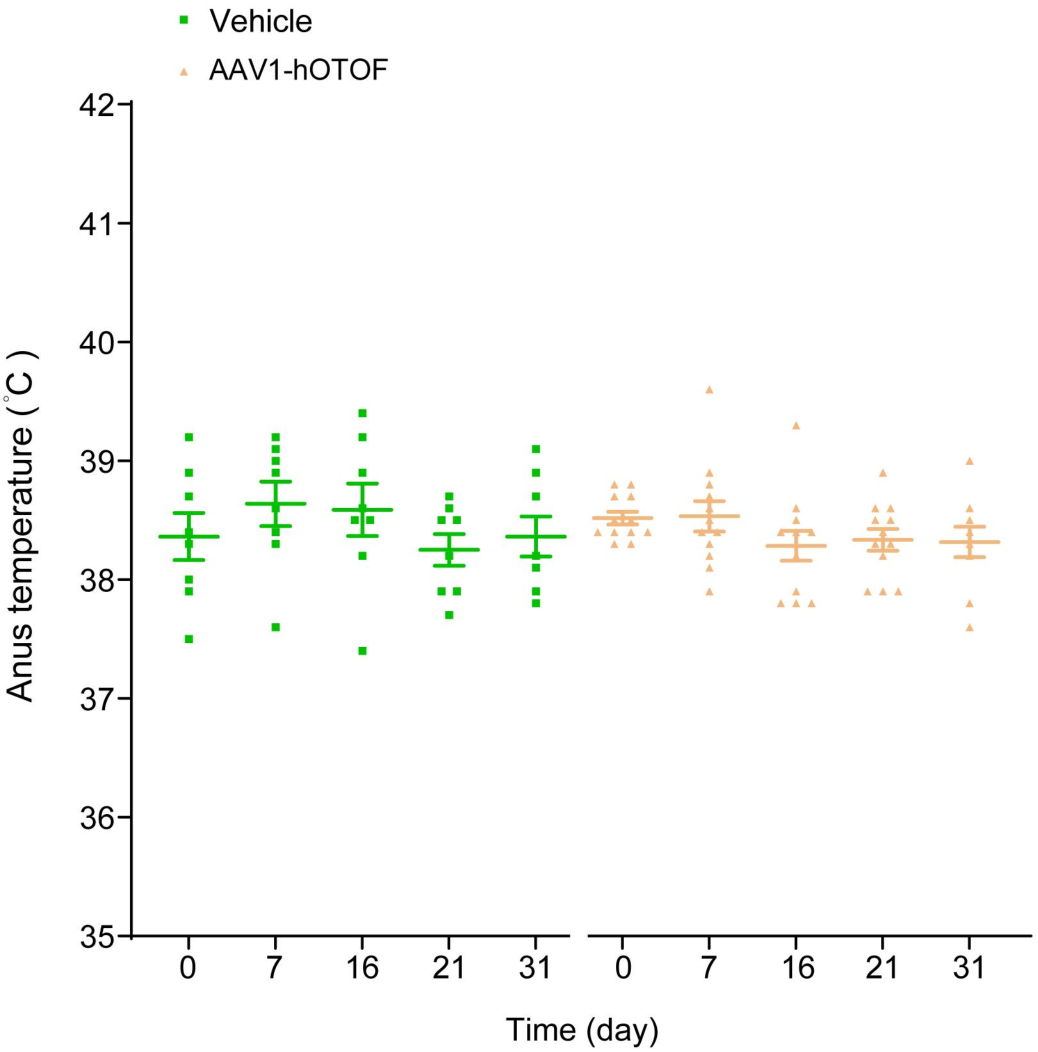


**Figure S5. Anus temperature in WT mice over time.** AAV1-hOTOF group, n = 12; vehicle group, n = 8. Data are displayed as mean ± SEM.

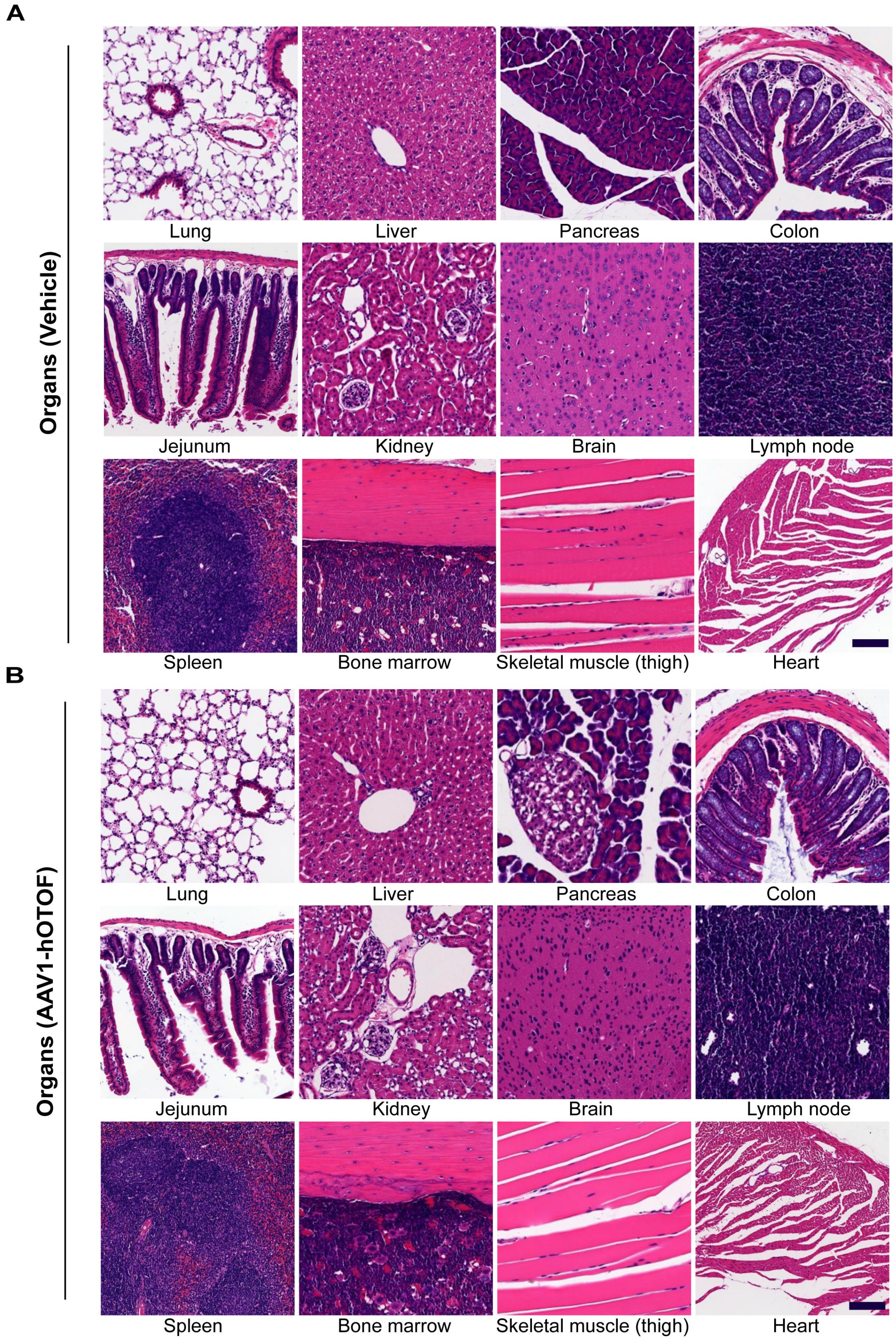


**Figure S6. Representative images of different organs from WT mice injected with AAV1-hOTOF** (**A**) **or vehicle** (**B**). No obvious signs of toxicity or inflammation were found. Tissues were stained with hematoxylin and eosin following standard procedures. Scale bar, 200 μm.

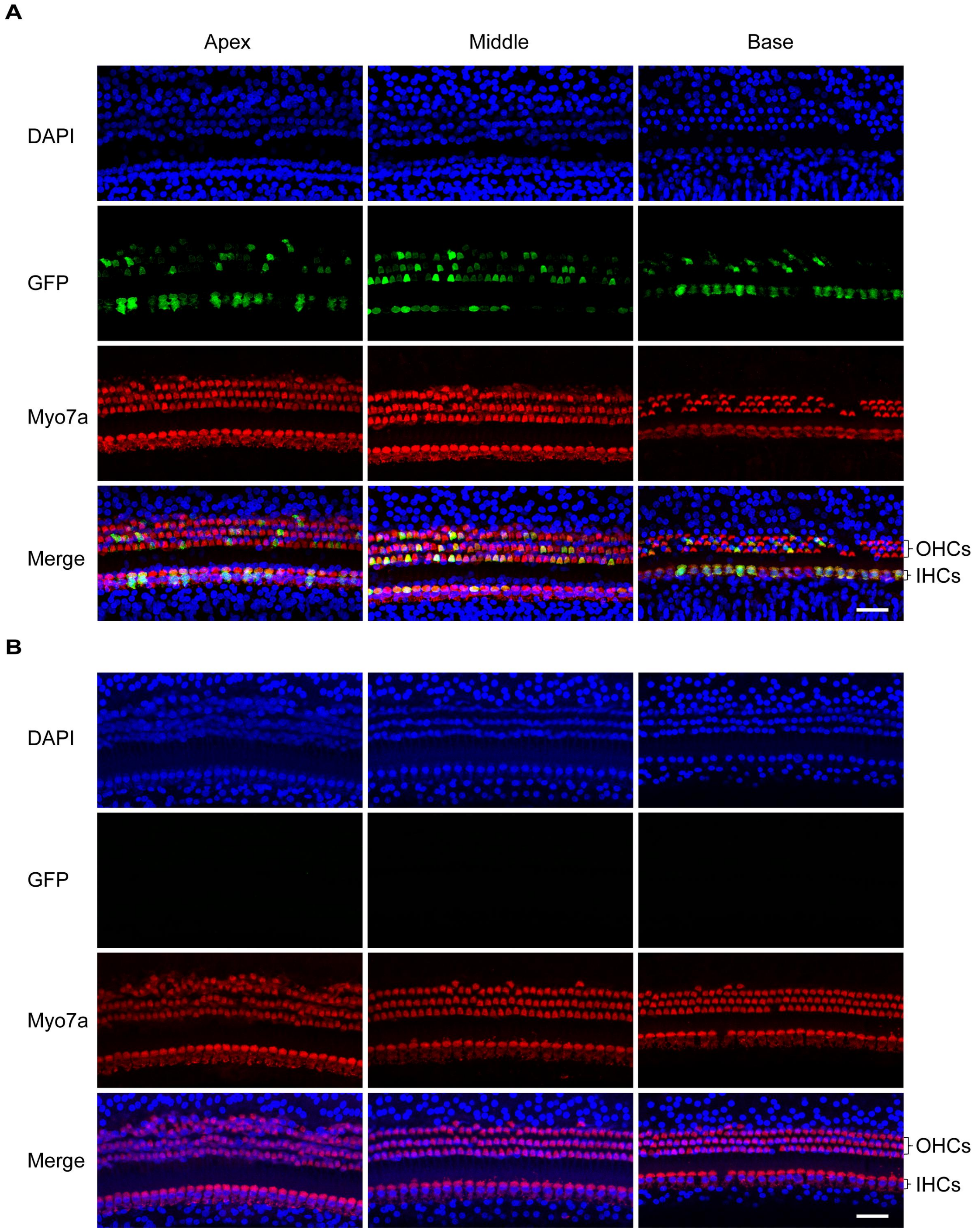


**Figure S7. Representative images of the expression of AAV1-GFP in NHP IHCs. A**. Representative images of Myo7a-labeled HCs in the injected ear of NHP at 4 weeks after injection with $1.5\times10^{11}$ vg AAV1-GFP. **B**. Representative images of Myo7a-labeled HCs in the contralateral uninjected ear of NHP after injection with $2.5\times10^{11}$ vg AAV1-GFP. No GFP signal was detected in the contralateral uninjected ear after injection. Scale bar, 20 μm.

**Table S1. Summary of the Irwin test recorded in WT mice injected with AAV1-hOTOF or vehicle**

| Treatment | | Incidence rate | | | | |
|---|---|---|---|---|---|---|
| | | Baseline | 1 Week | 2 Weeks | 3 Weeks | 4 Weeks |
| General behaviors | | | | | | |
| | Vehicle | 0/8 | 0/8 | 0/8 | 0/8 | 0/8 |
| | AAV1-hOTOF | 0/12 | 0/12 | 0/12 | 0/12 | 0/12 |
| Convulsive behaviors and excitability | | | | | | |
| | Vehicle | 0/8 | 0/8 | 0/8 | 0/8 | 0/8 |
| | AAV1-hOTOF | 0/12 | 0/12 | 0/12 | 0/12 | 0/12 |
| Reflex capabilities | | | | | | |
| | Vehicle | 0/8 | 0/8 | 0/8 | 0/8 | 0/8 |
| | AAV1-hOTOF | 0/12 | 0/12 | 0/12 | 0/8 | 0/12 |

(AAV1-hOTOF group, n = 12; vehicle group, n = 8).

**Table S2. The analyses of routine blood test in WT mice injected with AAV1-hOTOF or vehicle**

| Index | Vehicle | AAV1-hOTOF | *P* value |
|---|---|---|---|
| Mean red cell volume (fL) | 59.02 ± 0.65 | 58.68 ± 0.48 | 0.71 |
| Mean corpuscular hemoglobin concentration (g/dL) | 24.57 ± 0.29 | 24.80 ± 0.26 | 0.58 |
| Platelet count (×$10^3$ cells/μL) | 353.33 ± 17.61 | 350.13 ± 19.11 | 0.91 |
| White blood cell count (×$10^3$ cells/μL) | 1.84 ± 0.30 | 1.99 ± 0.31 | 0.74 |
| Red blood cell count (×$10^6$ cells/μL) | 3.84 ± 0.13 | 3.95 ± 0.21 | 0.64 |
| Hematocrit (%) | 22.58 ± 0.55 | 23.11 ± 1.13 | 0.64 |
| Mean platelet volume (fL) | 5.08 ± 0.09 | 5.09 ± 0.18 | 0.95 |
| Red blood cell distribution width (%) | 13.4 ± 0.13 | 13.21 ± 0.06 | 0.11 |
| Large uncountable cells (×$10^3$ cells/μL) | 0.02 ± 0.01 | 0.03 ± 0.02 | 0.66 |
| Mean corpuscular hemoglobin (pg) | 14.4 ± 0.06 | 14.56 ± 0.05 | 0.21 |
| Neutrophil (×$10^3$ cells/μL) | 0.53 ± 0.12 | 0.48 ± 0.19 | 0.81 |
| Lymphocyte (×$10^3$ cells/μL) | 1.21 ± 0.18 | 1.38 ± 0.15 | 0.50 |
| Monocyte (×$10^3$ cells/μL) | 0.03 ± 0.00 | 0.05 ± 0.12 | 0.16 |
| Eosinophil (×$10^3$ cells/μL) | 0.05 ± 0.01 | 0.05 ± 0.01 | 0.93 |
| Basophil (×$10^3$ cells/μL) | 0 | 0 | |
| Reticulocyte (×$10^9$ cells/μL) | 83.23 ± 13.05 | 93.73 ± 15.16 | 0.61 |
| Hemoglobin (g/dL) | 5.55 ± 0.19 | 5.74 ± 0.32 | 0.62 |

Data are displayed as mean ± SEM (AAV1-hOTOF group, n = 12; vehicle group, n = 8).

**Table S3. Serum chemistry values in WT mice injected with AAV1-hOTOF or vehicle**

| Index | Vehicle | AAV1-hOTOF | *P* value |
|---|---|---|---|
| Albumin (g/L) | 20.87 ± 0.31 | 21.82 ± 0.50 | 0.18 |
| Serum alanine aminotransferase (U/L) | 21.36 ± 1.61 | 28.80 ± 5.19 | 0.27 |
| Aspartate aminotransferase (U/L) | 150.25 ± 24.48 | 226.92 ± 52.99 | 0.28 |
| Total bile acid (μmol/L) | 1.63 ± 1.07 | 1.33 ± 0.43 | 0.78 |
| Total bilirubin (μmol/L) | 2.54 ± 0.23 | 2.26 ± 0.18 | 0.33 |
| Total protein (g/L) | 41.41 ± 0.60 | 44.01 ± 1.00 | 0.07 |

Data are displayed as mean ± SEM (AAV1-hOTOF group, n = 12; vehicle group, n = 8).

**Table S4. The information of the NHPs injected with AAV1-GFP**

| Animal ID | Age (year) | Gender | Weight (kg) | AAV serotype | Dose (vg) |
|---|---|---|---|---|---|
| #1 | 5.0 | male | 5.35 | AAV1 | $2.5\times10^{11}$ |
| #2 | 6.5 | male | 7.50 | AAV1 | $1.5\times10^{11}$ |

**Table S5. The analyses of routine blood test from NHPs injected with AAV1-GFP**

| Index | Days | 14 days | | 28 days | | Reference value |
|---|---|---|---|---|---|---|
| | Animal ID | #1 | #2 | #1 | #2 | |
| Mean red cell volume (fL) | | 81.90 | 76.90 | 81.70 | 77.00 | 77.70 ± 9.00 |
| Mean corpuscular hemoglobin concentration (g/L) | | 300.00 | 313.00 | 298.00 | 309.00 | 304.00 ± 26.00 |
| Platelet count (×10$^3$ cells/μL) | | 357.00 | 531.00 | 333.00 | 383.00 | 438.00 ± 184.00 |
| White blood cell count (×10$^3$ cells/μL) | | 9.52 | 13.08 | 8.44 | 9.59 | 11.42 ± 7.22 |
| Red blood cell count (×10$^6$ cells/μL) | | 4.98 | 5.22 | 5.07 | 5.37 | 5.89 ± 0.68 |
| Hematocrit (%) | | 40.80 | 40.20 | 41.40 | 41.40 | 45.70 ± 6.60 |
| Mean platelet volume (fL) | | 13.60 | 9.90 | 12.80 | 11.30 | 9.30 ± 2.40 |
| Red blood cell distribution width (%) | | 11.80 | 12.80 | 12.30 | 12.60 | 12.30 ± 1.40 |
| Mean corpuscular hemoglobin (pg) | | 24.60 | 24.10 | 24.30 | 23.80 | 23.60 ± 2.40 |
| Neutrophil (×10$^3$ cells/μL) | | 4.84 | 9.93 | 3.27 | 5.64 | 5.77 ± 7.14 |
| Lymphocyte (×10$^3$ cells/μL) | | 3.64 | 2.05 | 4.47 | 3.07 | 5.19 ± 4.16 |
| Monocyte (%) | | 8.90 | 7.30 | 6.60 | 7.30 | 2.70 ± 2.20 |
| Eosinophil (×10$^3$ cells/μL) | | 0.20 | 0.14 | 0.14 | 0.18 | 0.07 ± 0.16 |
| Basophil (×10$^3$ cells/μL) | | 0.00 | 0.00 | 0.00 | 0.00 | 0.04 ± 0.04 |
| Hemoglobin (g/dL) | | 122.00 | 126.00 | 123.00 | 128.00 | 139.00 ± 18.00 |
| Lymphocyte (%) | | 38.20 | 15.60 | 52.90 | 32.00 | 47.50 ± 31.20 |
| Neutrophil (%) | | 50.80 | 76.00 | 38.80 | 58.80 | 48.40 ± 32.80 |
| Eosinophil (%) | | 2.10 | 1.10 | 1.70 | 1.90 | 0.60 ± 1.20 |
| Basophil (%) | | 0.00 | 0.00 | 0.00 | 0.00 | 0.30 ± 0.40 |
| Monocyte (×10$^3$ cells/μL) | | 0.84 | 0.96 | 0.56 | 0.70 | 0.30 ± 0.28 |

Data are displayed as exact values. The values that were different from the normal values are highlighted in red (higher values) and blue (lower values).

**Table S6. The analyses of serum chemistry from NHPs injected with AAV1-GFP**

| Index | Days | 14 days | | 28 days | | Reference value |
|---|---|---|---|---|---|---|
| | Animal ID | #1 | #2 | #1 | #2 | |
| Albumin (g/L) | | 37.80 | 28.70 | 35.90 | 28.30 | 45.50 ± 4.60 |
| Alanine aminotransferase (U/L) | | 33.00 | 43.00 | 20.00 | 40.00 | 51.80 ± 48.00 |
| Aspartate aminotransferase (U/L) | | 16.00 | 20.00 | 18.00 | 22.00 | 61.40 ± 43.60 |
| Total bilirubin (μmol/L) | | 0.90 | 0.60 | 1.20 | 0.30 | 4.60 ± 3.76 |
| Total protein (g/L) | | 67.50 | 57.10 | 60.70 | 59.60 | 82.30 ± 11.60 |
| Creatinine (μmol/L) | | 83.00 | 46.00 | 71.00 | 42.00 | 83.10 ± 24.75 |
| Urea (mmol/L) | | 4.94 | 4.30 | 4.13 | 4.00 | 9.10 ± 3.28 |
| Glucose (mmol/L) | | 2.90 | 4.00 | 2.90 | 3.70 | 4.23 ± 3.06 |
| Total cholesterol (mmol/L) | | 3.14 | 3.00 | 2.63 | 3.53 | 3.89 ± 1.50 |
| Triglyceride (mmol/L) | | 0.74 | 0.83 | 0.38 | 0.34 | 0.43 ± 0.39 |
| Na (mmol/L) | | 145.00 | 145.00 | 136.00 | 144.00 | 151.00 ± 8.00 |
| P (mmol/L) | | 0.82 | 1.12 | 1.11 | 1.57 | 2.03 ± 0.91 |
| K (mmol/L) | | 4.60 | 4.20 | 4.30 | 4.10 | 5.21 ± 1.62 |
| Cl (mmol/L) | | 107.00 | 104.00 | 100.00 | 105.00 | 106.0 ± 3.00 |
| Ca (mmol/L) | | 2.45 | 2.32 | 2.30 | 2.29 | 2.66 ± 0.25 |
| Globulin (g/L) | | 29.70 | 28.40 | 24.80 | 31.30 | 36.50 ± 9.40 |
| Albumen/ globulin (Ratio) | | 1.30 | 1.00 | 1.40 | 0.90 | 1.25 ± 0.26 |

Data are displayed as exact values. The values that were different from the normal values are highlighted in red (higher values) and blue (lower values).
